# Midbrain activity supports high-level visual properties in primate temporal cortex

**DOI:** 10.1101/841155

**Authors:** Amarender R. Bogadhi, Leor N. Katz, Anil Bollimunta, David A. Leopold, Richard J. Krauzlis

**Author notes:** These authors contributed equally (Amarender R. Bogadhi, Leor N. Katz).

## Abstract

The evolution of the primate brain is marked by a dramatic increase in the number of neocortical areas that process visual information ^1^. This cortical expansion supports two hallmarks of high-level primate vision – the ability to selectively attend to particular visual features ^2^ and the ability to recognize a seemingly limitless number of complex visual objects ^3^. Given their prominent roles in high-level vision for primates, it is commonly assumed that these cortical processes supersede the earlier versions of these functions accomplished by the evolutionarily older brain structures that lie beneath the cortex. Contrary to this view, here we show that the superior colliculus (SC), a midbrain structure conserved across all vertebrates ^4^, is necessary for the normal expression of attention-related modulation and object selectivity in a newly identified region of macaque temporal cortex. Using a combination of psychophysics, causal perturbations and fMRI, we identified a localized region in the temporal cortex that is functionally dependent on the SC. Targeted electrophysiological recordings in this cortical region revealed neurons with strong attention-related modulation that was markedly reduced during attention deficits caused by SC inactivation. Many of these neurons also exhibited selectivity for particular visual objects, and this selectivity was also reduced during SC inactivation. Thus, the SC exerts a causal influence on high-level visual processing in cortex at a surprisingly late stage where attention and object selectivity converge, perhaps determined by the elemental forms of perceptual processing the SC has supported since before there was a neocortex.

## Main text

The primate superior colliculus (SC) in the midbrain is crucial for the normal control of visual attention, but whether and how SC activity interacts with cortical processing for attention is controversial ^5,6^. As a step toward identifying which areas of cortex might be influenced by the SC, we performed fMRI in monkeys performing attention tasks (figure 1a, b) and identified a novel attention-related region in the floor of the superior temporal sulcus (fSTS) of cortex (figure 1c; ^7^). By combining fMRI with and without SC inactivation, we found that the same fSTS region exhibited the largest reduction in attention-related BOLD modulation during the attention deficits caused by SC inactivation (figure 1d; ^8^). Relatively little is known about this cortical fSTS region ^9-14^ and it is unclear why it emerged in fMRI as the primary cortical area dependent on SC activity during the attention task. Our next step was to target this region for extracellular recording (figure 1e) to identify the signals conveyed by neurons and how these signals might depend on superior colliculus activity.

**Figure 1.**
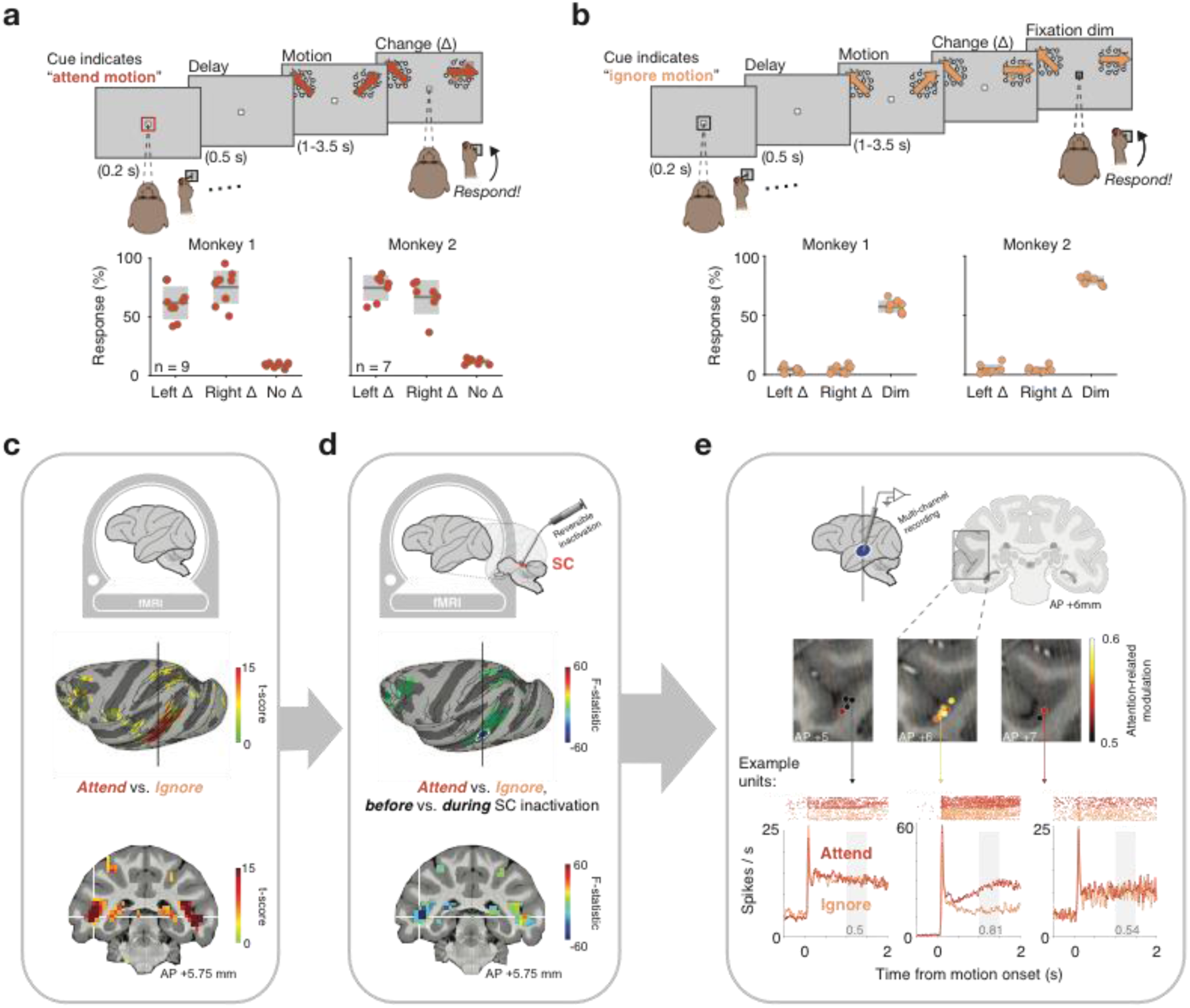
Identification of a cortical region dependent on SC activity. a. Top: Red cue instructed monkey to attend motion stimulus and report a change in motion direction (Δ). Bottom: response rates for left and right Δ (hits) and “no-Δ” (catch) trials. Individual sessions are plotted over gray box indicating mean ± sd. b. Top: Black cue instructed monkey to ignore motion-changes and report a dim in fixation spot. Bottom: response rates for left and right Δ (false alarms) and for dim (hits). Same format as a. c. Functional activations contrasting Attend and Ignore revealed maximal attention-related modulation in the fundus of STS (fSTS). d. Functional activations with and without SC inactivation identified a circumscribed region (blue colored patch) in fSTS whose modulation was the most dependent on SC activity. e. Coronal slices of the fSTS are overlaid with average attention modulation across neurons recorded in each location (colored spots). Oblique lines indicate electrode approach. Bottom: Example neuronal responses to *Attend* and *Ignore* conditions from three recording locations and their corresponding modulation values (gray text).

We recorded neuronal activity in the fSTS region, before and during SC inactivation, while two monkeys performed the same attention task used in fMRI experiments. In all conditions, monkeys maintained central fixation for the entirety of the trial and reported relevant stimulus changes by releasing a joystick. In the *Attend* condition (figure 1a), monkeys reported a change in the direction of motion of either of two peripheral visual stimuli. In the *Ignore* condition (figure 1b), monkeys ignored the behaviorally irrelevant direction changes in the peripheral stimuli and reported when the central fixation spot dimmed. The magnitude of stimulus change was adjusted such that the task was demanding and animals were at threshold performance – animals correctly responded in 73% ± 14% of trials in the *Attend* condition and 62% ± 13% in the *Ignore* condition (mean ± sd of both monkeys, see figure 1a, b for individual performance).

The targeted neuronal recordings during the attention task revealed a “hotspot” of attention-related modulation in fSTS (figure 1e). Neurons near the center of the fMRI-identified region tended to fire more in the *Attend* condition than in the *Ignore* condition compared to neurons in adjacent locations (figure 1e). Overall, fSTS neurons displayed strong attention-related modulation (figure 2a, c; Wilcoxon signed-rank test p < 6e-64, n = 380), and the median attention modulation index (AMI = 0.11; supplementary figure 1) was larger than what is typically observed in other areas of visual cortex (AMI = 0.051 in MT and 0.048 in MST ^15^).

**Figure 2.**
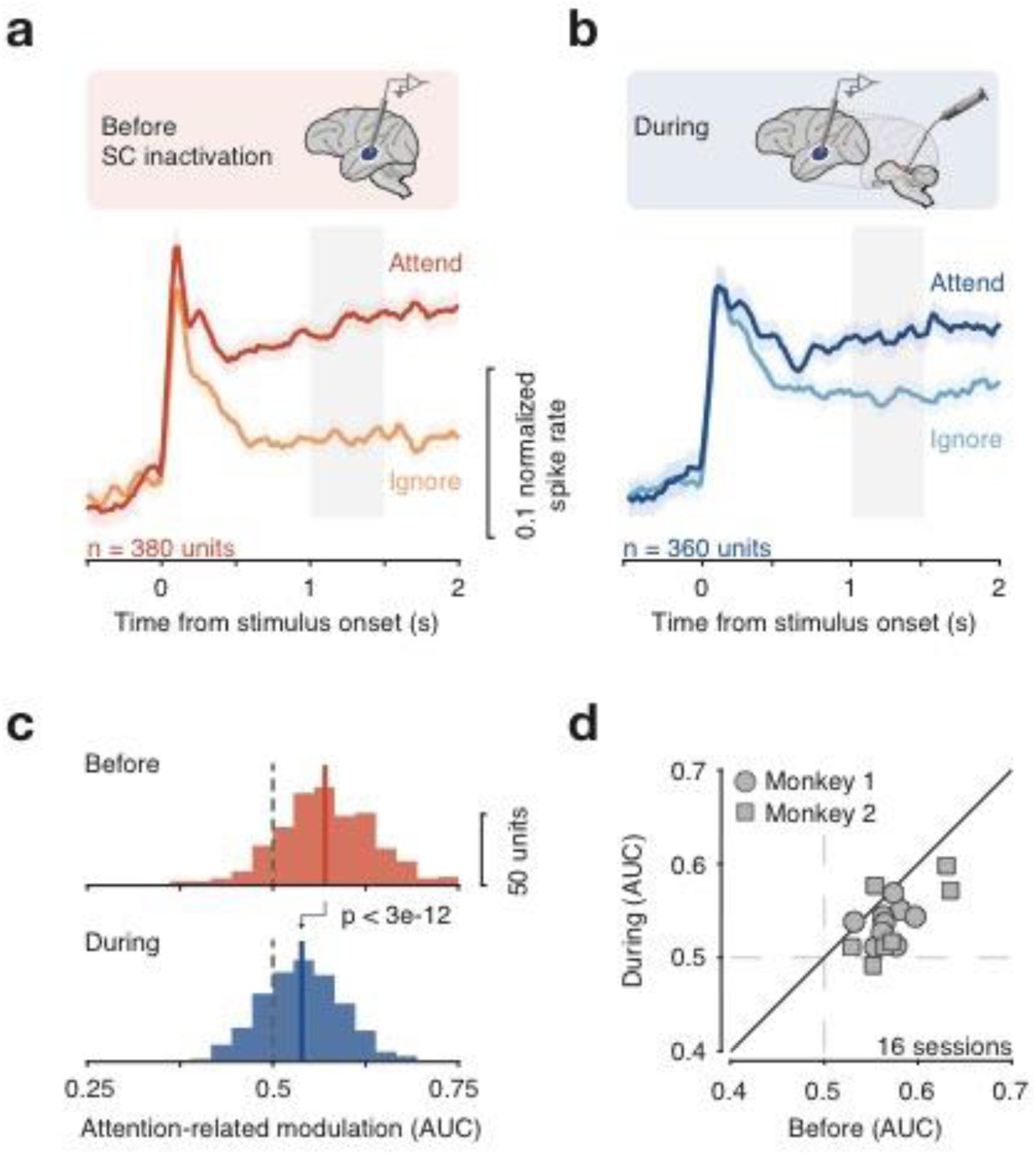
Attention-related modulation in fSTS neurons is dependent on SC activity. a-b. Population average of the normalized responses to motion stimulus in *Attend* and *Ignore* conditions before (a) and during (b) SC inactivation. Error bars indicate 68.2% CI. Grey window indicates time period used for computing attention-related modulation. c. Distribution of attention-related modulation values across neurons before and during SC inactivation. Solid and dotted lines indicate median and no modulation respectively. d. Average attention-related modulation across simultaneously recorded neurons within a session (mean ± sd = 23.75 ± 8.72), before and during SC inactivation.

We next tested whether this neuronal response modulation was affected by reversible inactivation of the SC. We confirmed that inactivation of the SC by microinjection of muscimol caused major deficits in attention task performance; consistent with previous findings ^16^, the deficits were spatially restricted to portions of the visual field contralateral to the injection site (supplementary figures 2-3). We found that the attentional modulation of fSTS neurons was markedly reduced by SC inactivation, unlike neurons in extrastriate visual cortex ^15^. The loss of attentional modulation was evident in the population averages of discharge rate over time (figure 2a, b), in individual neurons (figure 2c; Wilcoxon rank-sum test, p < 3e-12), across sessions (figure 2d; Wilcoxon signed-rank test, p < 0.002), and was not evident during control injections of saline (supplementary figure 4). Using a traditional attention modulation index (supplementary figure 1), modulation dropped by 45.9% during SC inactivation (from median of 0.11 before to 0.06 during). The reductions in attentional modulation cannot be explained by changes in neural spike rate or tuning properties since these were unchanged during SC inactivation (supplementary figure 5). These reductions also cannot be attributed to micro-saccades because the reductions in modulation were also observed in trials without micro-saccades (supplementary figure 6). These findings demonstrate that the large attention-related modulation of neurons in fSTS cortex depends on midbrain SC activity.

In addition to the attention-related modulation, neurons in fSTS also responded vigorously to the change in motion direction during the attention task (figure 3a). Since correct detection of the change was the crux of the attention task, this change-related activity is also an important feature of fSTS neurons. This change-related activity was also significantly attenuated during SC inactivation, as evident in the average population response to change (figure 3b), in single neurons (figure 3c; Wilcoxon rank-sum test, p < 4e-3), and across sessions (figure 3d; Wilcoxon signed-rank test, p < 0.03). We next quantified how well the trial-by-trial variation in change-related activity predicted the animal’s successful detections (a quantity termed *detect probability*). Before SC inactivation, detect probabilities were significantly greater than chance (figure 3g, Wilcoxon signed-rank test, p < 6e-64). During SC inactivation, detect probabilities were substantially reduced for both individual neurons (figure 3g; Wilcoxon rank-sum test, p < 3e-4) and experimental sessions (figure 3h; Wilcoxon signed-rank test, p < 0.03). Control saline injections again confirmed that these effects required suppression of SC activity (supplementary figure 7). These reductions in detect probability show that the degree to which fSTS activity can predict animal performance on a trial-by-trial basis also depends on SC activity.

**Figure 3.**
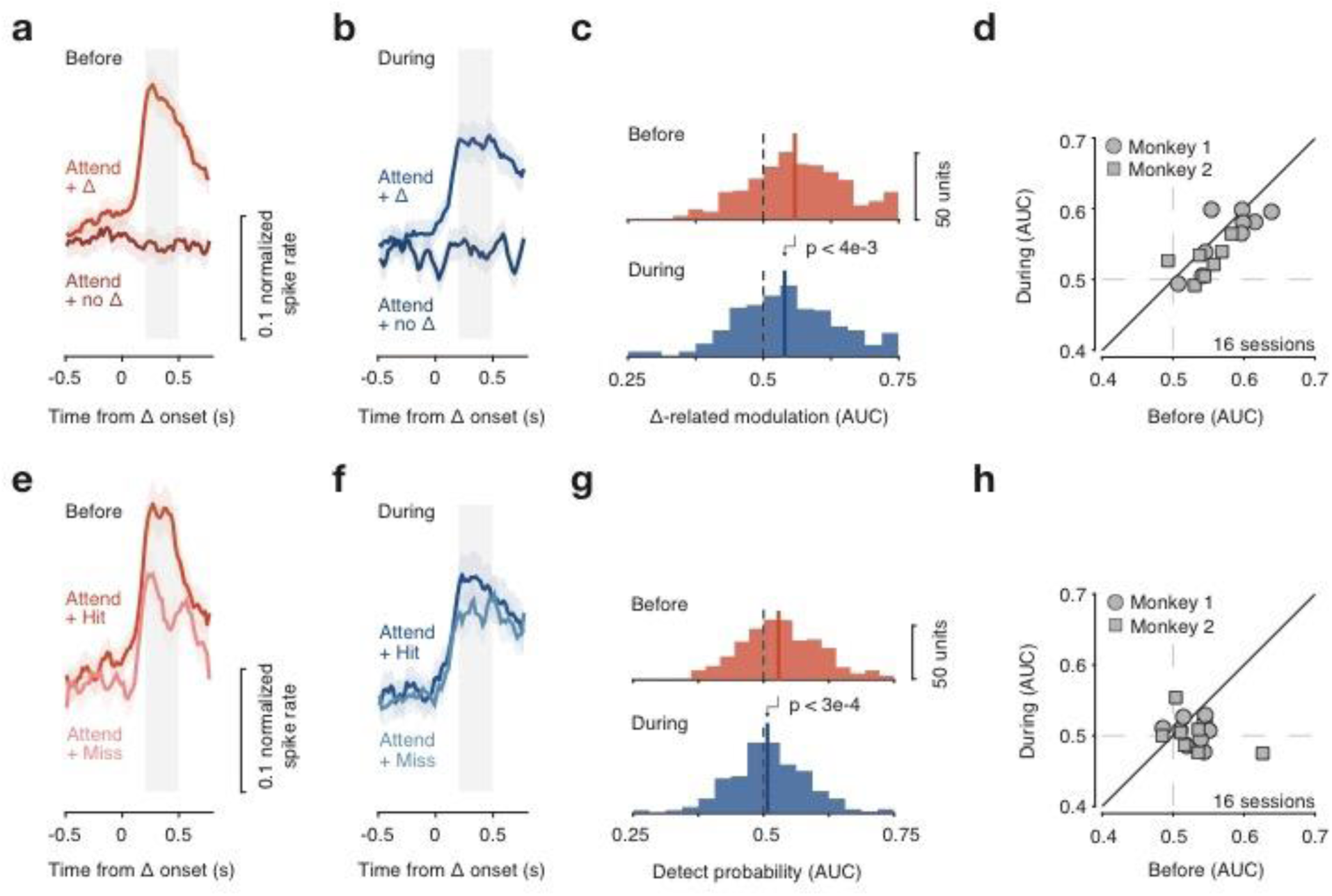
Change detection in fSTS neurons is dependent on SC activity. a-b. Population average of the normalized responses to motion-change (Δ) and no-change (no Δ) events in the *Attend* condition before (a) and during (b) SC inactivation. Grey window indicates time period used for computing change-related modulation. Error bars: 68.2% CI. c. Distribution of change-related modulation values before and during SC inactivation. Solid and dotted lines indicate median and no modulation respectively. d. Effect of SC inactivation on change-related modulation across sessions. e-h. Same conventions as in a-d but comparing hit and miss motion-change trials.

The loss of attention-related properties in fSTS neurons during SC inactivation was not specific to the feature of visual motion. Significant reductions in all of the neuronal measures reported – attention-related modulation, change-related modulation, and detect probabilities – were also observed (supplementary figure 8) when the experiments were repeated using a different visual stimulus (a brief pulse of second-order orientation) that was designed such that it cannot be detected using motion energy ^8^.

These results identify the fSTS region as part of a novel circuit for attention control that depends on SC activity. The properties of this control mechanism are different from the well-known mechanisms centered in the prefrontal cortex, which are thought to provide feedback signals to early visual areas that regulate how basic visual features are processed ^17^. First, fSTS lies at a later stage in the hierarchy of cortical visual areas ^18,19^. Second, the control of fSTS region by the SC is not specific to a particular visual feature (supplementary figure 8). Third, the changes in fSTS caused by SC inactivation omitted two signatures of selective attention typically found in earlier visual areas – neither spike-count correlations nor Fano factor ^20,21^ showed consistent effects during the attention deficits caused by reversible inactivation (supplementary figure 9).

Given the proximity of fSTS to visual form processing areas and the lack of feature specificity in its dependence on SC, we speculated that the SC control of fSTS region might be related to elemental perceptual forms and events rather than the detailed representation of visual features afforded by the expansion of neocortex. To this end, we tested for another visual property that is common in the temporal lobe but usually studied separately from attention – selectivity for visual objects ^22-24^. Monkeys passively viewed a stream of briefly (250 ms) flashed images of visual objects within the receptive fields of neurons recorded in the fSTS. To control for low-level visual features, we also included grid- and phase-scrambled versions of the same images randomly intermixed in the stream (figure 4a).

**Figure 4.**
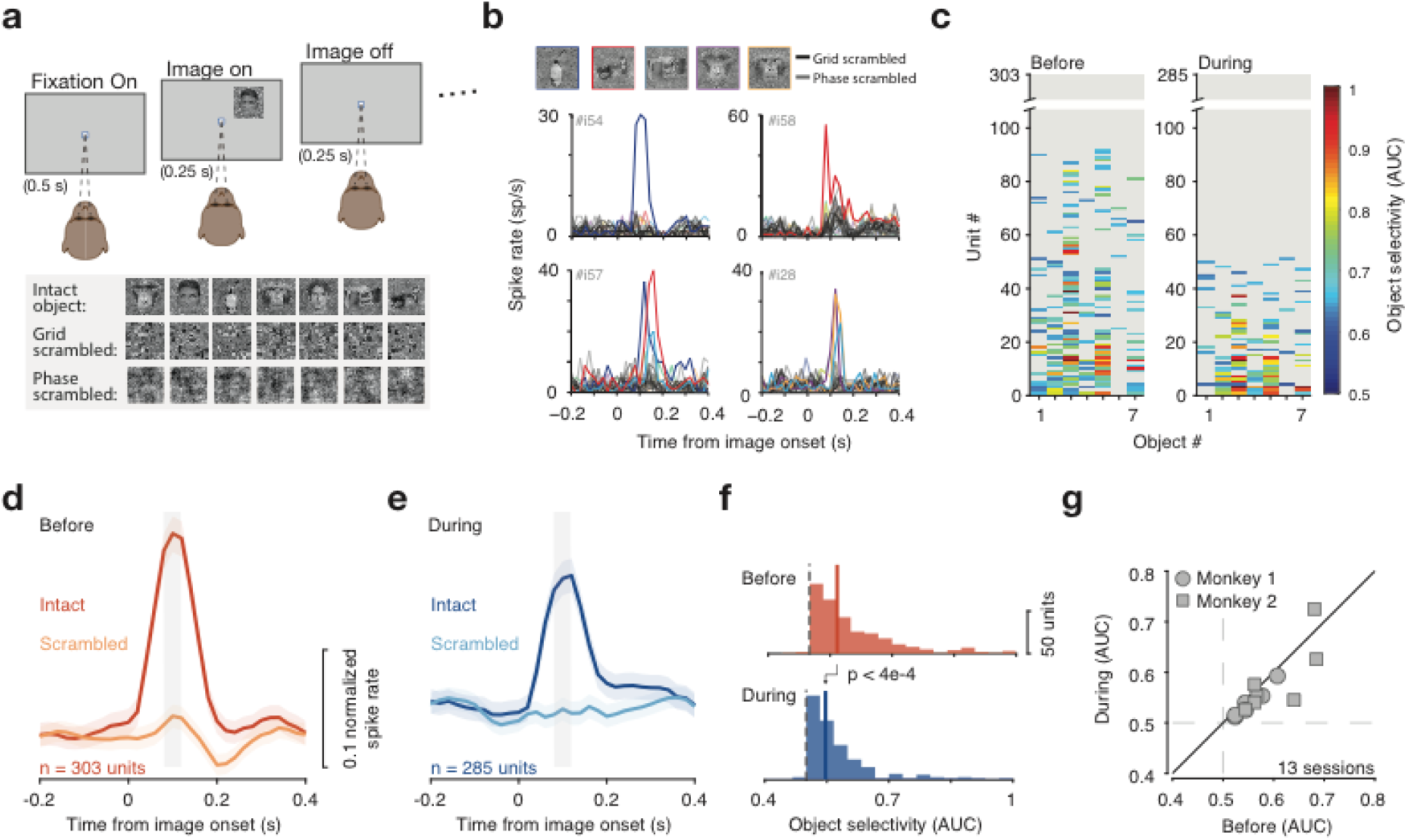
Object selectivity in fSTS neurons is dependent on SC activity. a. Monkeys fixated a central spot while either object, grid-scrambled, or phase-scrambled images were presented. b. Example neurons that responded selectively to individual object images. c. Object selectivity values for individual neurons, sorted by number of objects selective for, before and during SC inactivation. Non-significant selectivity (p > 0.05) is shown as grey. d-e. Population average of normalized responses to the most selective object and the corresponding scrambled object, before (d) and during (e) SC inactivation. Grey window indicates time period used for computing object selectivity. Error bars: 68.2% CI. f. Distribution of object selectivity values before and during SC inactivation. Solid and dotted lines indicate median and no selectivity respectively. g. Effect of SC inactivation on object-selectivity across sessions.

About one-third of the neurons we recorded exhibited strong selectivity for particular visual objects. For example, neurons responded vigorously during the presentation of a particular image (e.g. “water bottle”), but not during any of the other intact or scrambled images (figure 4b). Across the population of 303 neurons, 92 showed significant selectivity for at least one object (figure 4c), and this selectivity cannot be attributed to attention drawn to particular images, because neurons recorded at the same time showed selectivity for different images (e.g. units #54, #57 & #58 in figure 4b). The rapid rise, strength and prevalence of object selectivity was evident in the population responses (figure 4d) and the distribution of object selectivity values (median = 0.56, significantly greater than 0.5, Wilcoxon signed-rank test, p < 2e-51).

During SC inactivation, the number of object-selective neurons was substantially reduced from 30.3% (92/303) to 17.89% (51/285) (figure 4c; Chi-square test, p < 7e-4). Significant reductions in object-selectivity during SC inactivation were evident in population responses (figure 4d, e) and individual neurons (Figure 4f; Wilcoxon rank-sum test, p < 4e-4). The reduction in object-selectivity was consistent across sessions (Figure 4h; Wilcoxon signed-rank test, p < 0.04) and monkeys (supplementary figure 10). This finding demonstrates that fSTS neurons gain their basic visual object selectivity, in part, through a pathway that involves the superior colliculus.

In summary, our findings demonstrate that the primate SC plays a causal role in the cortical processing of visual stimuli through a previously undistinguished patch of cortex in the fSTS. SC activity probably reaches fSTS through the thalamic pulvinar ^25-27^. Similar ascending circuits from SC through higher-order thalamus to later stages of visual cortex have been recently described in mice ^28-30^. The functions of these circuits from the SC may complement those of the well-known hierarchy of visual pathways starting in the lateral geniculate nucleus of the thalamus ^18,19^. We suggest that the cortical visual system might use signals from the SC as a shortcut for rapid object processing: rather than relying on feedforward hierarchical circuits to identify all possible objects in a scene, inputs from the SC might guide the efficient recognition of objects at the specific location deemed most important at that moment.

## Methods

### Animals

Two adult male rhesus monkeys (*Macaca mulatta*) weighing 7-9 kg were used in the study. All procedures and animal care were approved by the National Eye Institute Animal Care and Use Committee and complied with the Public Health Service Policy on the humane care and use of laboratory animals. We surgically implanted plastic head-posts and electrophysiology chambers, under isoflurane and aseptic conditions, to access the SC and the fundus of the STS (fSTS). The placement of the fSTS chamber was guided by previously obtained functional imaging results ^8^, and we targeted the fSTS region in different hemispheres across the two monkeys. Both monkeys were trained on an attention task (figure 1a, b), a visually guided saccade task (for mapping the affected region during SC inactivation, supplementary figure 2), an object tuning task (figure 4a), and several other passive fixation tasks (for mapping spatial receptive fields and direction tuning of single neurons).

### Experimental apparatus

Animals were seated and head-fixed in a primate chair (Crist Instrument Inc., Hagerstown, MD, and custom-built) inside a darkened booth at a distance of 48 cm from a VIEWPixx display (VPixx Technologies, Saint-Bruno, QC Canada) with a 1920 × 1200 resolution (∼ 60° × 38°) and 100Hz frame-rate. Stimuli were presented on the VIEWPixx display that was controlled by a mid-2010 Mac Pro (Apple Inc., Cupertino, CA) running MATLAB (The Mathworks, Natick, MA) with the Psychophysics Toolbox extensions ^32^. The background luminance of the screen across all tasks was 14 cd/m^2^. Eye position was recorded using an EyeLink 1000 infrared eye-tracking system (SR Research Ltd., Ottowa, Ontario, Canada); this signal was monitored online to ensure strict fixation within a 2° fixation window during all tasks. The primate chairs were equipped with a single axis joystick (CH Products, model HFX-10) that the monkeys used to report relevant stimulus changes during the attention task. Joystick presses and releases were detected by a change in voltage signal. All experiments were controlled using a modified version of PLDAPS ^33^.

### Electrophysiology recording

Neuronal and behavioural signals (e.g. eye position, joystick) were acquired by an Omniplex system (Plexon Inc., Dallas, Texas). Neuronal signals were recorded using a 24-channel linear array (V-probes, Plexon Inc., Dallas, Texas) that was introduced into the fSTS region by a custom built motorized micro-drive. Figure 1e shows coronal sections of fSTS region overlaid with colored spots representing average attention-related modulation (see below) of all neurons recorded in that grid hole. AP and ML location of the colored spots were determined by the grid hole, depth was determined as the average depths across recording sessions in that grid hole. In additional sessions, we recorded a few millimeters away from the ‘hotspot’ and found either non-standard responses to the stimulus onset or no response at all (supplementary figure 11), indicating that our region of interest in the fSTS is circumscribed.

Entry into the target fSTS region was confirmed based on depth measurements compared to the MRI slices (figure 1e), and changes in neural activity observed during transitions from gray matter in the dorsal bank of the STS to sulcus, and from sulcus to gray matter in the target region of the fSTS. Upon entry into the target region we allowed the V-probe to settle and the neuronal activity to stabilize (∼ 1 hour) before beginning the experimental session.

Each session began with an online mapping of spatial receptive fields to optimize stimulus placement for the subsequent tuning and attention tasks. Following the mapping, we collected neuronal responses during the direction tuning, attention, and object tuning tasks. At the end of the “before SC inactivation” set of tasks, we reversibly inactivated SC, and continued collecting data for the “during SC inactivation” set of tasks, starting with the attention task, followed by the tuning and receptive field mapping tasks. Overall, we performed 22 experimental sessions with SC inactivation: 16 with motion-change detection stimulus (9 in monkey #1; 7 in monkey #2); 6 with orientation pulse detection stimulus (in monkey #1), and 8 additional control sessions of saline injection and sham with the motion-change detection stimulus (monkey #1).

### Receptive field mapping task

Spatial receptive fields of fSTS neurons were mapped in each experimental session before and during SC inactivation, while monkeys performed a passive fixation task. Monkeys started each trial by fixating a central square spot (0.21°wide; green colour). Following 0.5 s of fixation, a random dot motion stimulus was presented briefly (0.25 s on and 0.25 s off) within a circular aperture (3° radius) at 100% coherence. The stimulus was presented at locations drawn randomly from an XY grid (5° spacing, covering the display) and at locations selected by the experimenter based on the neuronal responses. The direction of the motion stimulus was pseudo-randomized across stimulus presentations. Each trial lasted for 4 seconds and consisted of 6 stimulus presentations. Monkeys were required to maintain fixation within a 2° window for the full duration of the trial to receive a reward. We collected an average of 552 (before) and 522 (during) presentations in each experimental session.

After receptive field mapping, a location was selected by the experimenter which maximized the spatial overlap of receptive fields across the recording channels. This selected location was thereafter fixed for all subsequent tasks: the direction tuning task, the object tuning task, and the attention task, before and during SC inactivation. Overall, locations used ranged between 8 – 14° eccentricity across all sessions.

### Direction tuning task

Direction tuning properties of fSTS neurons were mapped in each experimental session before and during SC inactivation, while animals performed a passive fixation task. The fixation task was similar to the receptive field mapping task, except that each brief motion stimulus presentation (0.25 s on and 0.25 s off) was presented at the fixed visual field location. Random dot motion was presented at 100% coherence and the direction of motion was selected pseudo-randomly on each stimulus presentation from one of the 12 directions (equally spaced from 0 to 360 degrees). A 0% coherence stimulus was also included. Each trial lasted for 4 seconds and consisted of 6 stimulus presentations. Overall, we collected an average of 40 (before) and 36 (during) presentations per motion direction in each experimental session.

### Object tuning task

Monkeys performed a passive fixation task similar to the direction tuning task, except that the motion stimulus was replaced with static images containing objects (visual forms) and their corresponding grid- and Fourier phase-scrambled versions (figure 4a). A total of 21 unique images were used (7 object, 7 grid-scrambled and 7 phase-scrambled images) and all object images were corrected for luminance histogram and Fourier spectra using SHINE toolbox ^34^, before generating their corresponding scrambled versions. Each image was presented briefly (0.25 s on and 0.25 s off) at the fixed location selected previously. Each trial lasted 4 seconds and consisted of 6 image presentations. Overall, we collected an average of 49 (before) and 40 (during) presentations per image in each experimental session.

### Attention task

The attention task consisted of two main conditions: *Attend* and *Ignore* (figure 1a, b), plus two control conditions, *Attend Single Patch* and *Baseline*. In all task conditions, monkeys initiated the trial by pressing the joystick down and fixating a central square spot (50 cd/m^2^). Following 0.3 s of fixation, a cue in the form of a colored square outline was flashed around the central fixation spot for 0.2 s. The width of the fixation spot was 0.21°and the width of the cue was 0.35°. The color of the cue instructed the monkeys to attend or ignore the peripheral motion stimuli and detect the relevant events. In the *Attend* and *Attend Single Patch* trials, the color of the central cue was red, indicating that the relevant event was a direction-change in the peripheral motion stimulus (detailed below). In the *Ignore* and *Baseline* trials, the color of the central cue was black, indicating that direction-change in the peripheral stimulus was irrelevant and the luminance change in the fixation spot was relevant. In all conditions, monkeys were required to maintain central fixation within a 2° fixation window for the full duration of the trial. All four task conditions were presented in a block design where each condition was presented once (in pseudo-random order) before advancing to the next block. Each block consisted of 150 trials: 54 *Attend*; 54 *Ignore;* 30 *Attend Single Patch;* and 12 *Baseline*. Monkeys completed an average of 5 blocks before SC inactivation and 4 blocks during inactivation across 16 sessions. The sequences of events in each condition are presented below.

In *Attend* trials, following 0.5 s after cue offset, two random-dot motion stimuli appeared on either side of the fixation spot in the contralateral and ipsilateral hemi-fields, symmetric to the vertical meridian. The location of the contralateral stimulus matched the location from mapping tasks (see Receptive field mapping section). Overall, location eccentricity ranged between 8 – 14° across sessions. On 77.7% of *Attend* trials, one of the motion stimuli changed direction during 1-3.5 s after stimulus onset (uniform probability), and the monkeys responded by releasing the joystick within 0.3 – 0.8 s to receive a reward. On the remaining 22.3% trials, there was no stimulus change and monkeys were rewarded for continuing to hold the joystick pressed (catch trials).

In *Ignore* trials, the peripheral motion stimuli and task sequence were identical to those in the *Attend* trials, but the monkeys were rewarded for ignoring the peripheral stimuli, and instead, responded to a change in fixation spot luminance (decrease of 1-2 cd/m^2^ across sessions). On 77.7% of *Ignore* trials, the fixation spot luminance change occurred during 1–3.5 s after stimulus onset (uniform probability) and monkeys responded by releasing the joystick within 0.3 – 0.6 s to receive a reward. On the remaining 22.3% of trials, the fixation spot luminance remained unchanged, and monkeys were rewarded for continuing to hold the joystick pressed (“catch trials”). Independent of the luminance change, on 66.6% of the trials, one of the two motion stimuli changed direction and the monkeys were rewarded for ignoring the motion direction-change and responding to the change in the fixation spot luminance.

*Attend Single Patch* trials were identical to *Attend* trials, with only one of the two motion stimuli presented on any given trial. The stimulus was presented either at the same contralateral or ipsilateral location on equal number of trials.

*Baseline* trials were identical to *Ignore* trials, except that no peripheral stimulus was presented. In this condition, monkeys were simply required to detect a change in fixation spot luminance.

### Random dot motion stimulus

Each random dot motion stimulus consisted of moving dots presented in a circular 3° radius aperture. The motion direction of each dot was drawn from a normal distribution with a mean value of 30° above horizontal (in all but one session) and a standard deviation of 16°, to be consistent with the previously performed fMRI study ^8^. The lifetime (10 frames, 100 ms), density (25 dots/^o2^/s), and speed of the dots (15 °/s) were held constant. Luminance of each moving dot in the motion patches was 50 cd/m^2^. The change in direction of motion (Δ) ranged from 0.75 to 1.25 standard deviations of mean motion direction across monkeys and sessions.

### Attention tasks: Orientation-pulse detection

In separate sessions, monkey 1 performed a modified version of attention task with the same conditions (*Attend, Ignore, Attend Single Patch, Baseline*), where random dot motion stimulus was replaced with dynamic white noise stimulus and motion-change event was replaced with a brief (0.5 s) second-order orientation pulse. In the *Attend* and *Attend Single Patch* conditions, the monkey reported the orientation pulse event by releasing the joystick within 0.3 - 0.8 s to get a reward, whereas in the *Ignore* condition, monkey ignored the orientation pulse event and reported the luminance decrease in the fixation spot by releasing the joystick within 0.3 - 0.6 s to get a reward. We collected an average of 6 blocks before SC inactivation and 5 blocks during SC inactivation across 6 sessions.

The white noise stimulus had a diameter of 6°and consisted of checks, each the size of a pixel, dynamically changing luminance on every frame at random, with luminance values ranging from 8 to 84 cd/m^2^. The second-order orientation stimulus was generated by briefly (0.5 s) modulating the contrast of a white noise stimulus with a 2-dimensional sinusoid, whose spatial frequency and orientation was 0.7 cycles/deg and 90°respectively. Importantly, the mean luminance of the stimulus (38 cd/m^2^) was held constant throughout its presentation and was constant across every band in the oriented grating. We refer to this as a second-order orientation stimulus, because the visibility of the oriented grating was due to local differences in contrast, not luminance, and we confirmed that the orientation pulse contained no change in motion energy ^8^.

### Reversible inactivation

Reversible inactivation of the intermediate layers of the SC (n = 22; monkey #1 (left hemisphere): 15, monkey #2 (right hemisphere): 7) was done by injecting muscimol (0.3-0.5 μl; 5 mg/ml) based on methods described previously ^16^. All candidate sites were first identified by neuronal recordings and electrical stimulation prior to the muscimol inactivation experiment. Injections were done using a custom-made apparatus ^35^ using an injection pump (Legato, KD Scientific) at a constant rate of 0.05 μl/min. For saline controls, the same volumes were injected at the same locations in the intermediate layers of the SC at the same rates.

### Mapping of the impaired visual field

Thirty minutes after the end of the muscimol injection, we used a visually guided saccade task to map the affected visual field using methods described previously ^16^. We measured the peak velocity for saccades made to targets at different visual field locations before and during inactivation. The locations at which saccade peak velocities were reduced indicated the locus and the extent of the affected visual field (supplementary figure 2).

### Electrophysiology analysis

Electrophysiological data obtained from the linear arrays were sorted offline into single neurons using the *kilosort* algorithm followed by a manual curation in the *phy* software ^36^. For analyses related to attention tasks, we included single neurons with an average trial firing rate greater than 1 spike/s in both *Attend* and *Ignore* blocks. The trial firing rate of a neuron was computed as total spikes over the full duration of trial divided by the trial duration. This resulted in inclusion of 380/563 and 360/529 for the before and during conditions, respectively.

For each neuron, mean peri-stimulus time histogram (PSTH) was computed by binning spike times in 10ms non-overlapping bins (figure 1e, figure 4b). To compute the average population response in different tasks and epochs (figures 2, 3, 4), we normalized the mean PSTH of each neuron by its peak activity at the stimulus onset in the corresponding task and averaged the normalized PSTHs across all units, followed by a smoothing operation with a 5-point moving average.

#### Attention-related modulation

We used spike counts in the *Attend* and *Ignore* conditions during the delay period (1 – 1.5 s after stimulus onset, grey window in figure 2a, b) as signal and noise respectively to compute the area under the receiver operator characteristic curve (AUC). This AUC value determined a neuron’s attention-related modulation. To evaluate the effect of SC inactivation on attention-related modulation we computed the distributions of AUC values before and during SC inactivation (figure 2c). We also computed the traditional attention-related modulation index (AMI) as 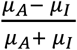, where *μ*_*A*_ is the average spike count during the delay period (1 – 1.5 s after stimulus onset) in *Attend* condition and *μ*_*I*_ is the average spike count in *Ignore* condition. The effects reported in this manuscript do not depend on whether we use our attention-related modulation AUC measure or AMI (supplementary figure 1).

#### Fano factor index and neural correlations

In the same epoch as attention-related modulation (1 – 1.5 s after stimulus onset, grey window in figure 2a, b), we computed fano-factor index and neuronal correlation index for each neuron. Fano-factor index was computed as 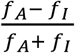, where *f*_*A*_ is the fano-factor during *Attend* and *f*_*I*_ is the fano-factor during *Ignore*; *f*_*A*_ was computed as 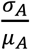, where *σ*_*A*_ was the variance in spike count during *Attend* condition (supplementary figure 9). Neuronal correlation index was computed as the difference in inter-neuronal spike count correlations during *Attend* and *Ignore*. Spike count correlation between a pair of neurons was computed as a Pearson’s correlation coefficient (MATLAB). Overall, we computed correlations for 4856 pairs of neurons before SC inactivation and 4341 pairs during SC inactivation (supplementary figure 9).

#### Change-related modulation

To compute an AUC value that determined a neuron’s change-related modulation in activity we used spike counts in the *Attend* condition for change trials and no-change trials, during the post-change period (0.2 – 0.5 s epoch following the change, grey window in figure 3a, b).

#### Detect probability

To compute an AUC value for each neuron’s detect probability ^37^ we compared spike counts in the *Attend* condition for change trials, when the monkey successfully detected a stimulus change (“hit”) to when the monkey failed to detect a stimulus change (“miss”), during the post-change period (0.2 – 0.5 s epoch following the change, grey window in figure 3e, f).

#### Object selectivity

To compute an AUC value for each neuron’s object selectivity in the object-tuning task, we used spike counts for the object image compared to the corresponding scrambled version of the image during the 0.08 – 0.12 s epoch after the image onset (grey window in figure 4d, e).

## Data Availability

The datasets generated during and/or analysed during the current study are available from the corresponding author on reasonable request.

## Acknowledgments

We thank Tom Ruffner and Nick Nichols for technical support. We are grateful to Fabrice Arcizet, James Herman, and Lupeng Wang for helpful discussions and input. This work was supported by the National Eye Institute Intramural Research Program at the National Institutes of Health.

## Author Contributions

All authors designed the experiments. A.R.B., L.N.K., and A.B. conducted experiments and collected the data. A.R.B. and L.N.K analyzed the data. All authors interpreted the results. R.J.K., A.R.B. and L.N.K. wrote the manuscript.

---

**Supplementary Figure 1.**
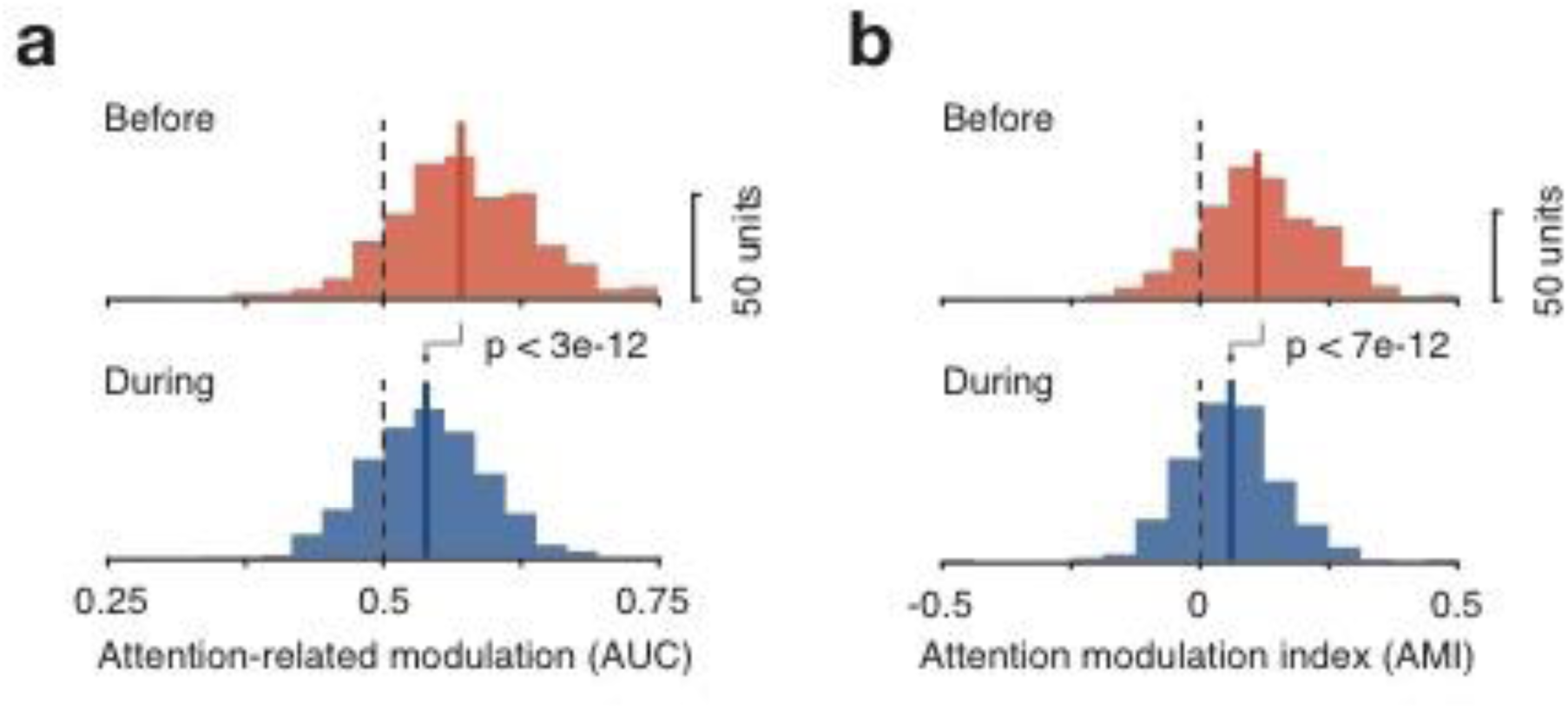
Effect of SC inactivation on attention-related modulation in fSTS is consistent across different methods of quantification. a. Distribution of area under the receiver operator curve (AUC) values quantifying attention-related modulation (see Methods) before (top) and during SC inactivation (bottom) (Same as figure 2c). b. Distribution of attention modulation index (AMI) values quantifying attention-related modulation (see Methods) before (top) and during SC inactivation (bottom). The difference in distributions is highly significant (see p-values on plot) and similar in both the AUC and AMI cases.

**Supplementary Figure 2.**
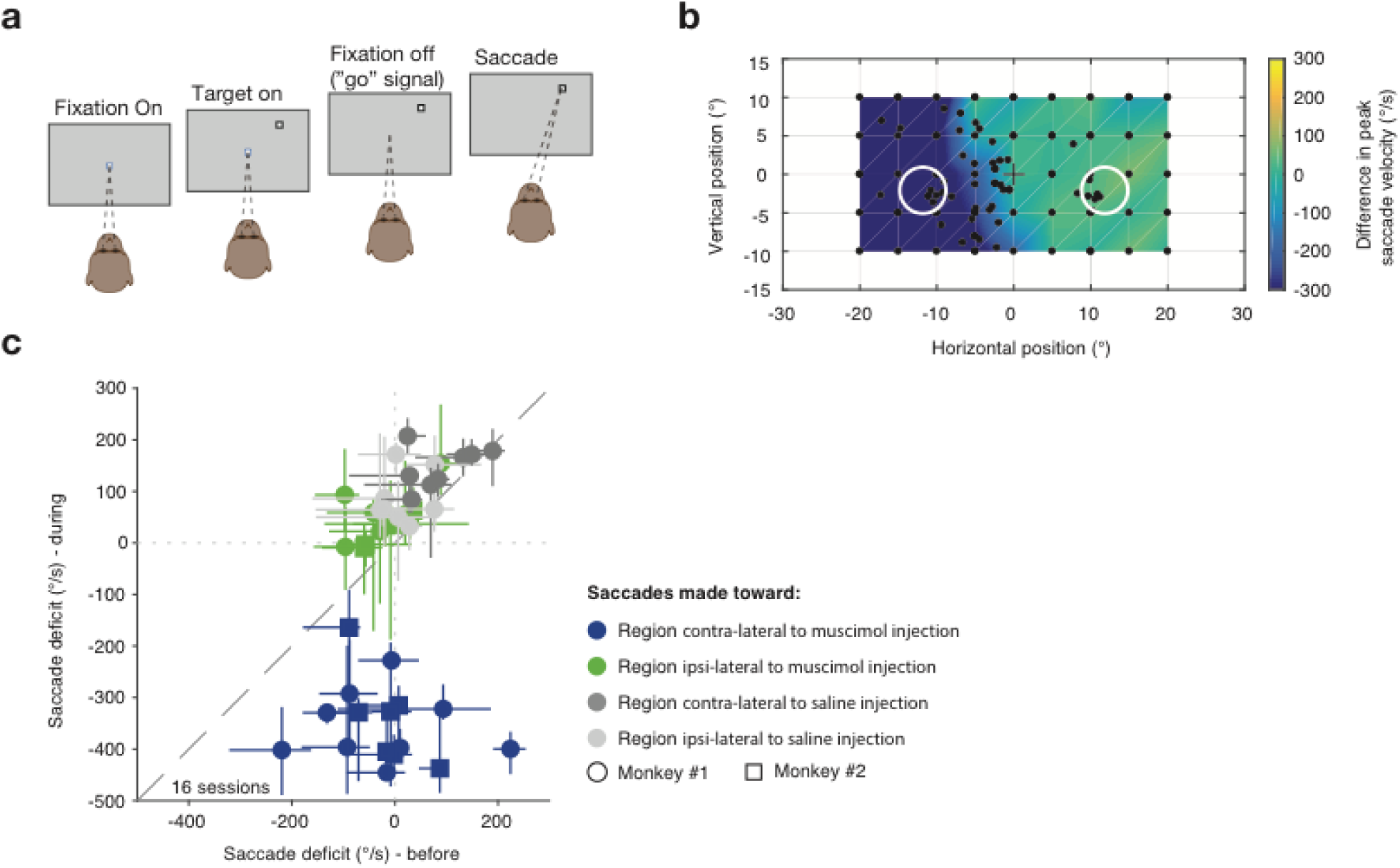
SC inactivation was confirmed using a visually guided saccade task. a. Sequence of events in the visually guided saccade task. Following fixation of a central spot, a saccade target was presented at a visual field location. The subject is rewarded for successfully making a saccade to the target upon fixation offset (see methods). b. A visual field map of saccadic deficit induced by SC inactivation in an example session. The difference between during and before SC inactivation saccade peak velocities is plotted for a large number of sampled targets in space. White circles represent the location of the motion stimulus used in the attention task that followed the visually guided saccade task. c. Summary of saccade deficit in the contralateral and ipsilateral stimulus locations (white circles in b) before and during injection for muscimol (SC inactivation) and saline injection sessions.

**Supplementary Figure 3.**
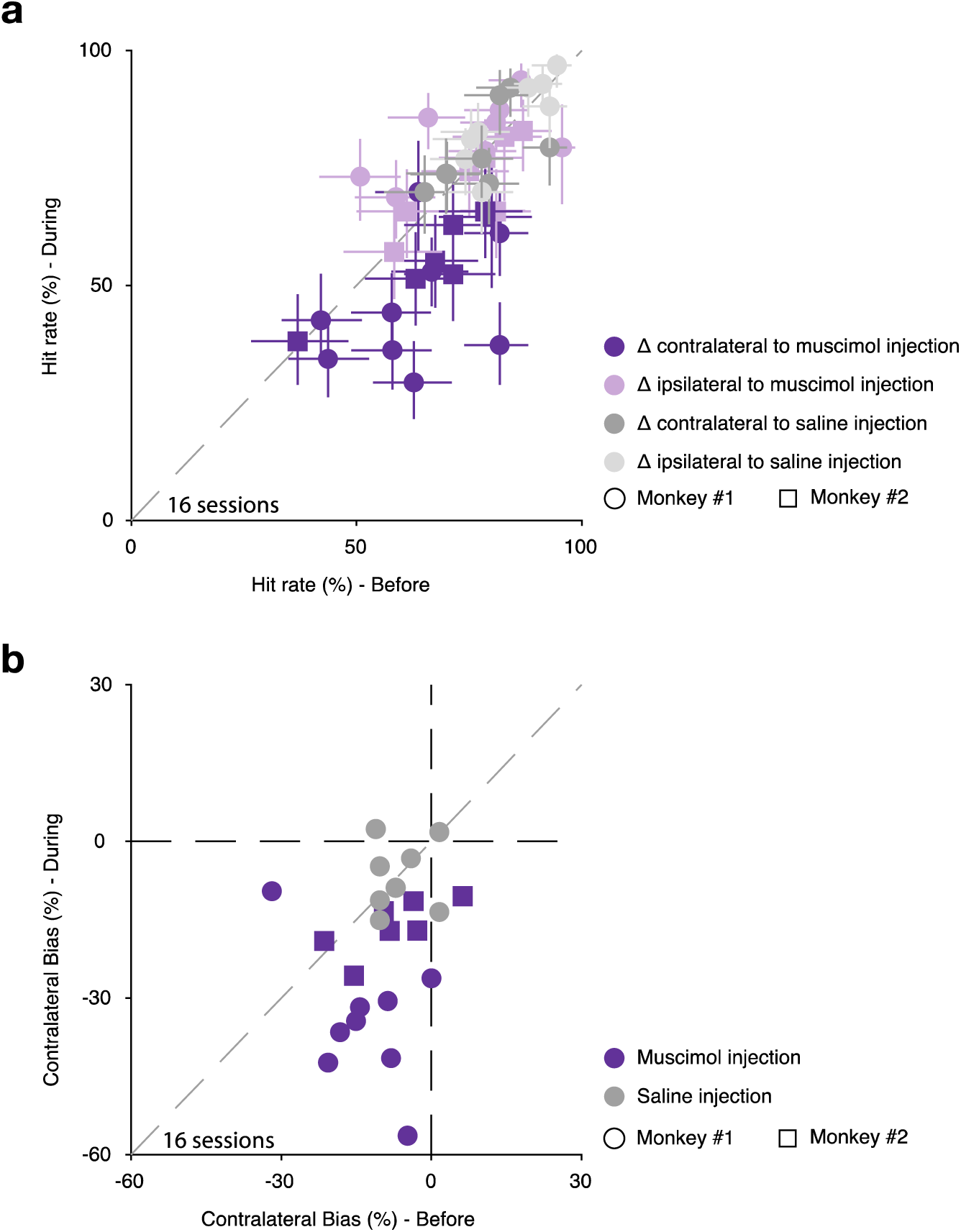
Behavioral deficits in the attention task during SC inactivation. The percentage of correctly detected motion-direction changes (hit rate) in the contralateral (affected) visual hemifield was significantly reduced during SC inactivation compared to performance in the ipsilateral (unaffected) visual hemifield (Wilcoxon signed-rank test, p < 5e-04) and to performance in the contralateral hemifield before inactivation (Wilcoxon signed-rank test, p < 0.002). a. The successful detection of a change (Δ) in motion direction (hit rate) in the *Attend* condition before SC inactivation vs. during, for when the change occurred in either the contralateral or ipsilateral stimulus, across all muscimol and saline injection sessions. b. The difference in hit rate between contralateral and ipsilateral motion direction changes in the *Attend* condition (termed “contralateral bias”), before SC inactivation vs. during, across all muscimol and saline injection sessions (same data as in a).

**Supplementary Figure 4.**
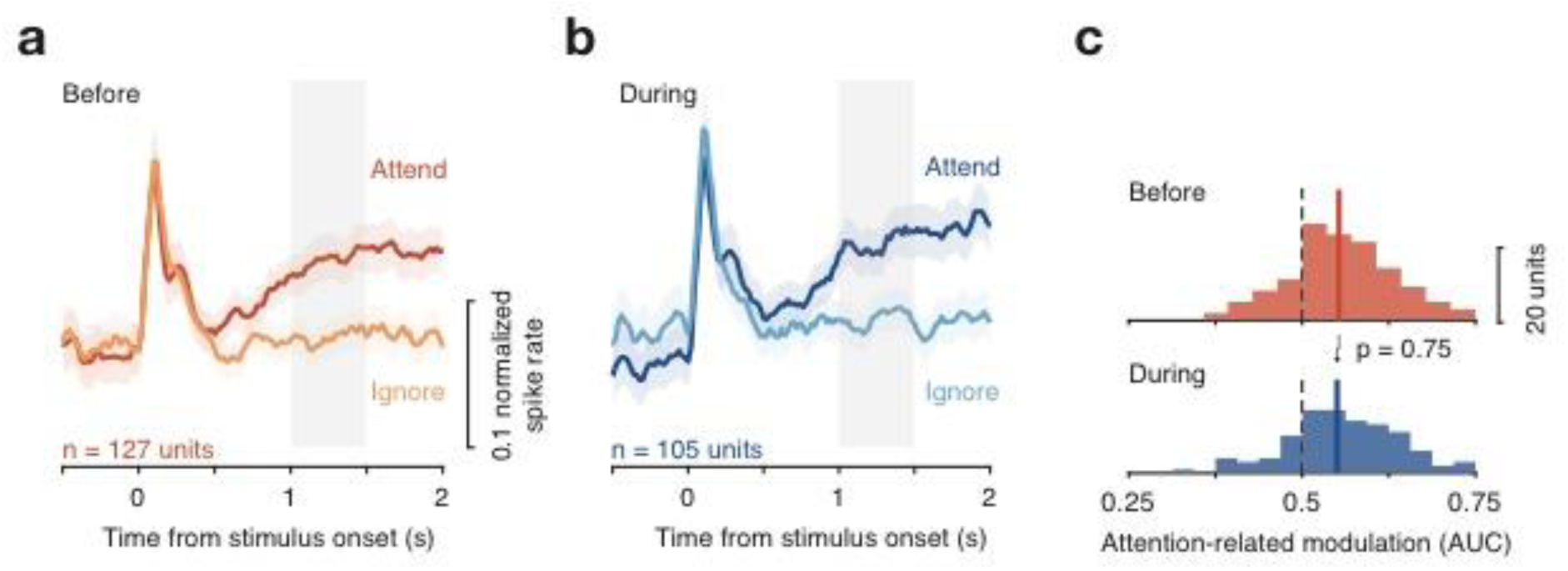
Saline and sham injections had no effect on attention-related modulation in fSTS. a-c. Attention-related modulation before and during saline/sham injections (same format as figure 2a-c).

**Supplementary Figure 5.**
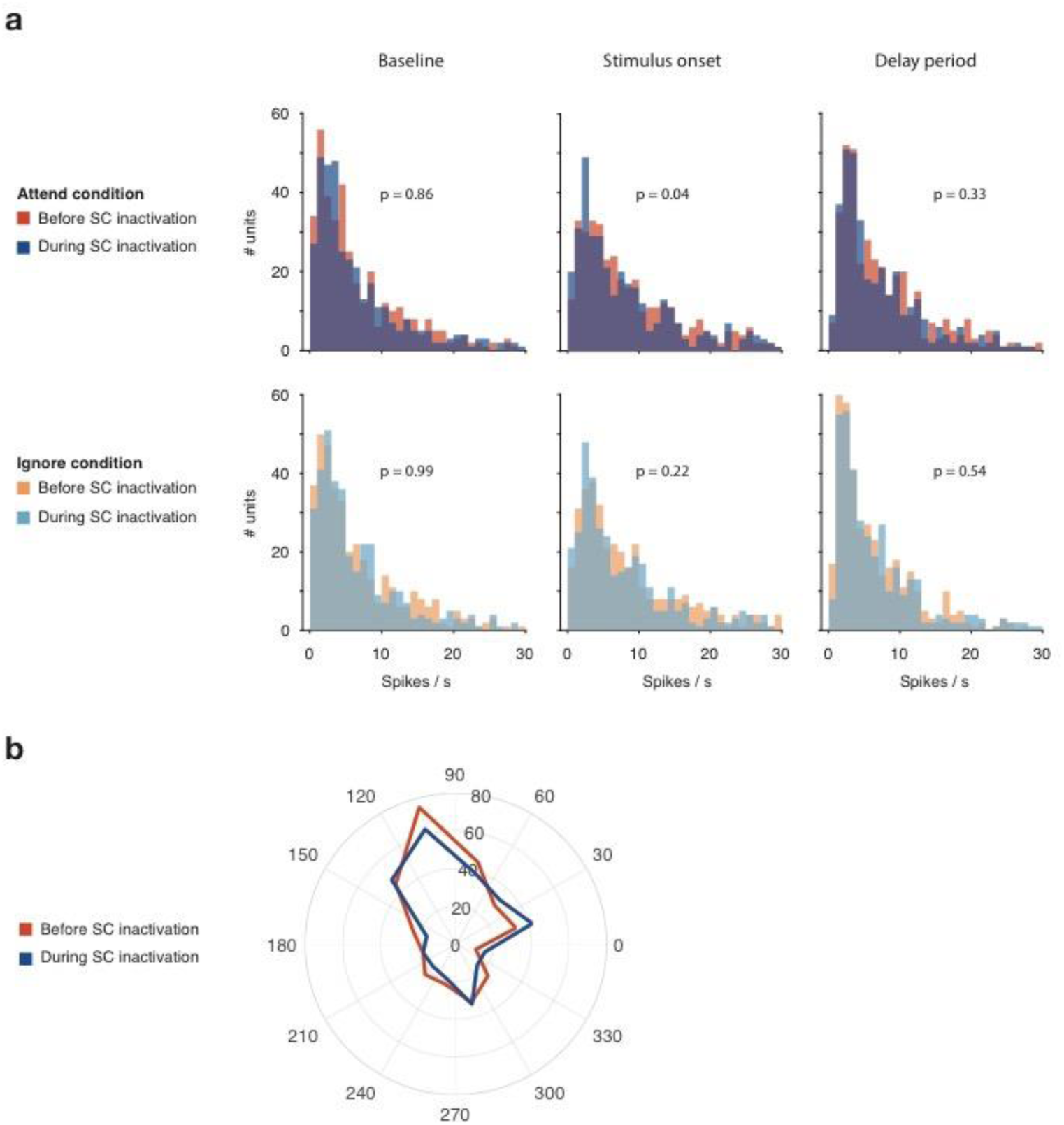
SC inactivation did not affect neuronal firing rate or direction tuning. a. Spike rate was computed in three key epochs of *Attend* and *Ignore* conditions: baseline (0.12-0.8s before stimulus onset); stimulus onset (0.8-0.12s after stimulus onset); delay period (1-1.5s after stimulus onset). The distributions of spike rates before and during SC inactivation are shown in the three epochs (up to 30sp/s for visualization) of *Attend* (top) and *Ignore* (bottom) conditions. p-values comparing median firing rate before and during SC inactivation (Wilcoxon rank-sum test) are noted on individual panels. b. Preferred direction of each neuron was obtained using the direction tuning task (see Methods). Histogram of preferred directions across the fSTS population of neurons, before and during SC inactivation, is plotted in polar coordinates. Radial measure represents number of neurons with the corresponding preferred direction.

**Supplementary Figure 6.**
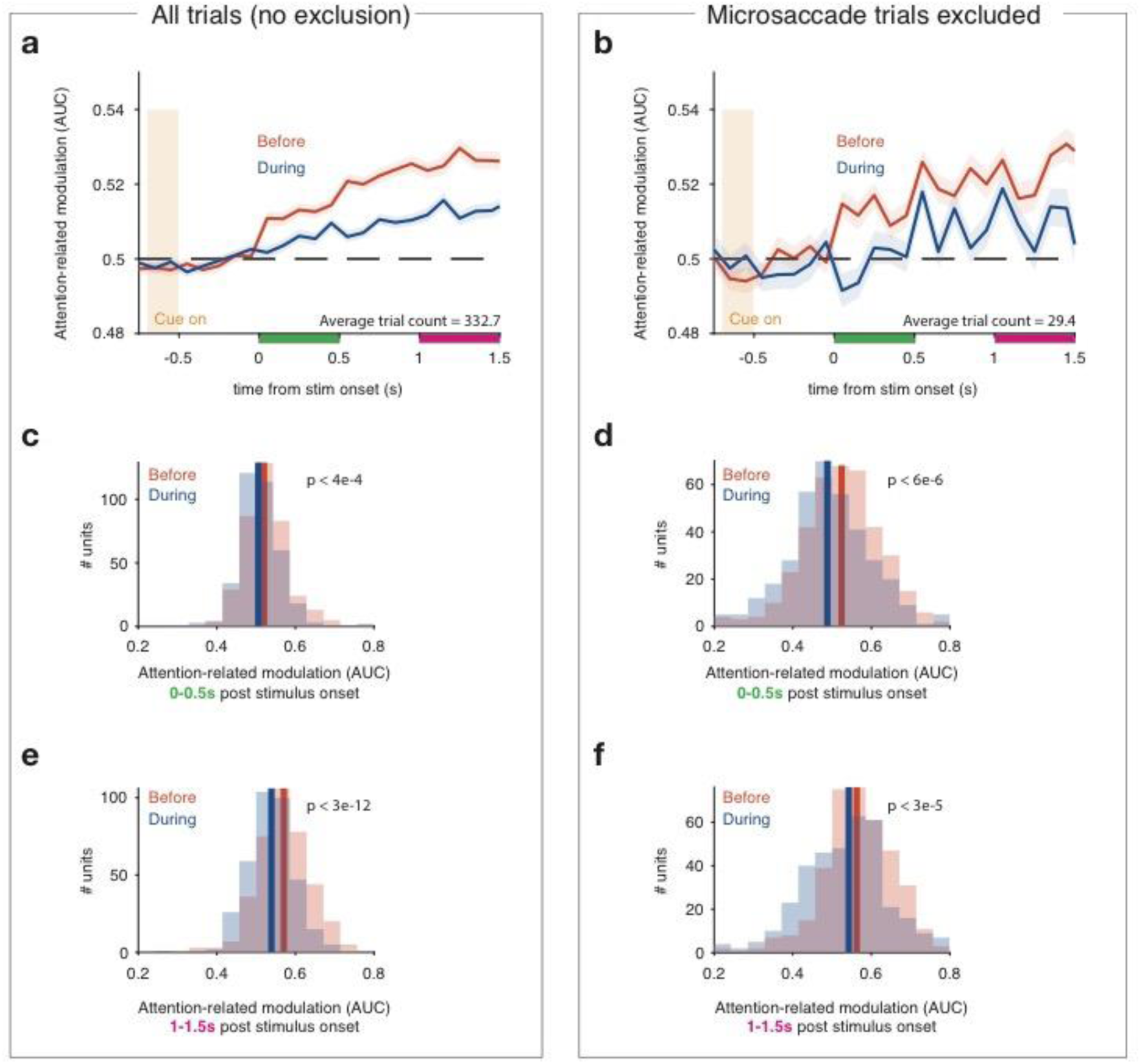
Effect of SC inactivation on attention-related modulation in fSTS cannot be attributed to microsaccades. A recent study ^31^ reported that attention-related modulation in visual cortex occurs only following a microsaccade in the direction of the cued location. To determine the influence microsaccades might have had on our results, we detected all microsaccades during the full duration of a trial (amplitude: mean ± sd = 0.42° ± 0.2°) and excluded trials in which a microsaccade occurred anywhere from cue onset to the end of the time window in which we measured attention-related modulation (1-1.5s post stimulus onset; figure 2). Excluding trials with microsaccades anywhere from cue onset until the end of this 1-1.5s window (magenta box under x-axis in a, b) resulted in too few trials per neuron to compute an AUC (mean ± sd = 4.83 ± 5.85). The earliest time window that could be used for this analysis while still yielding sufficient trials for statistical power was the 0-0.5s window aligned to stimulus onset (green box under x-axis in a, b). By measuring attention-related modulation in this 0-0.5s window on trials with no microsaccade from cue onset until the end of this 0-0.5s time window, before and during SC inactivation, we show that the effect of SC inactivation on attention-related modulation in fSTS cannot be attributed to microsaccades. a-b. Time-course of attention-related modulation before and during SC inactivation, computed on all trials (a) and on trials with no microsaccade from cue onset to 0.5s post stimulus onset (b). Even without microsaccades from cue onset to 0.5s post stimulus (panel b), the attention-related modulation is significant following stimulus onset, and is reduced during SC inactivation. c-d. Distribution of attention-related modulation values before and during SC inactivation, computed in the 0-0.5s time window (green box under x-axis in a, b), on all trials (c) and trials with no microsaccade from cue onset to 0.5s post stimulus onset (d). Solid lines indicate medians. The results in d demonstrate that even with no microsaccades preceding the observed modulation, neurons in fSTS show attention-related modulation that is dependent on SC activity. e-f. Same format as c-d, but attention-related modulation was computed in the standard time window (1-1.5s, magenta box under x-axis in a, b) using the same trials as in c, d. We present these data in order to directly relate this analysis with the analyses reported in the main text, which uses the 1-1.5s window (figure 2c), but note that it is still possible that microsaccades occurred after our exclusion window (−0.7 to 0.5s relative to stimulus onset) and before the delay period (1-1.5s post stimulus onset).

**Supplementary Figure 7.**
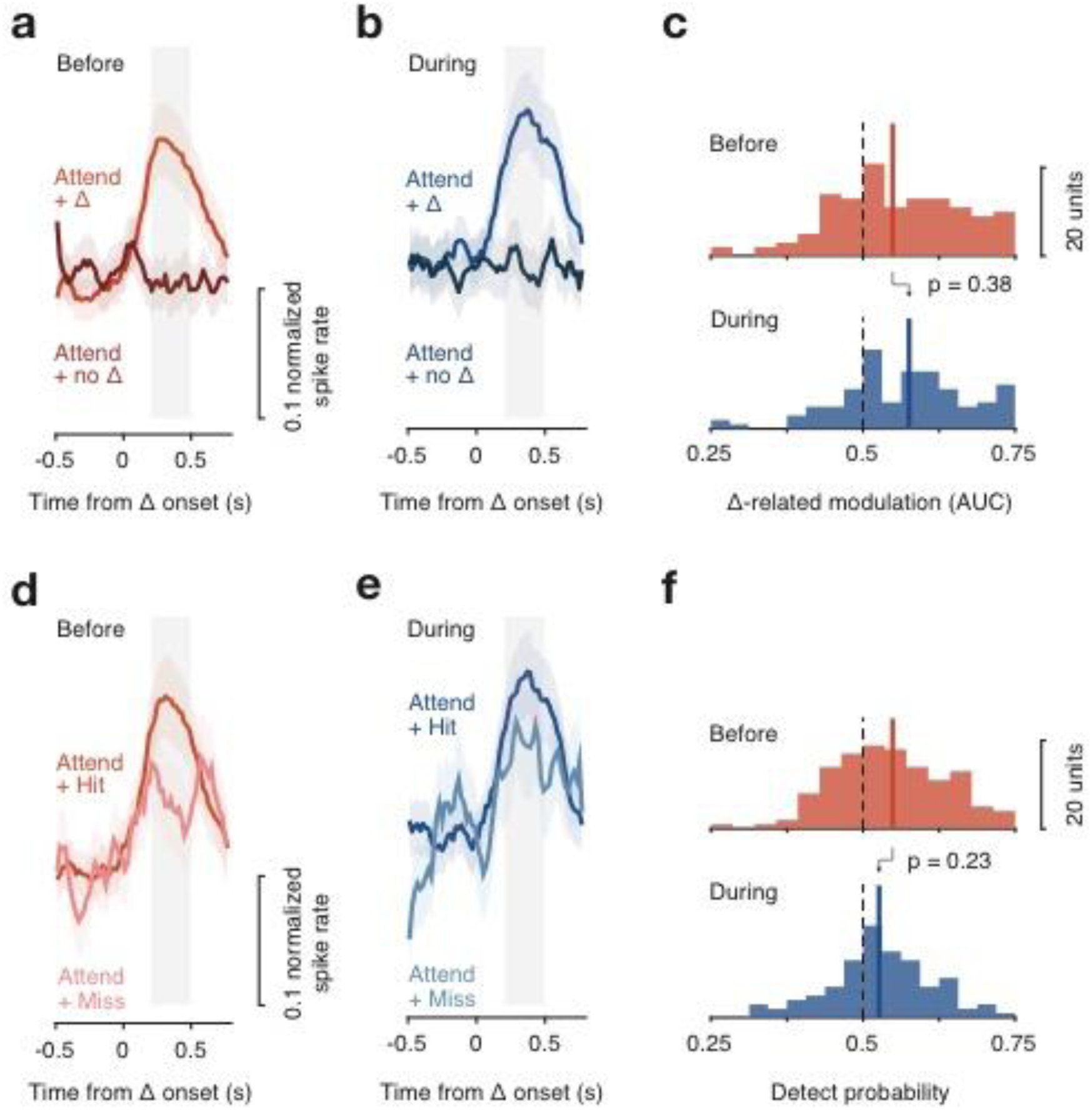
Saline and sham injections had no effect on change-related activity or detect probability in fSTS. a-c. Change related activity before and during saline/sham injections (same format as Figure 3a-c). d-f. Detect probability before and during saline/sham injections (same format as Figure 3e-g).

**Supplementary Figure 8.**
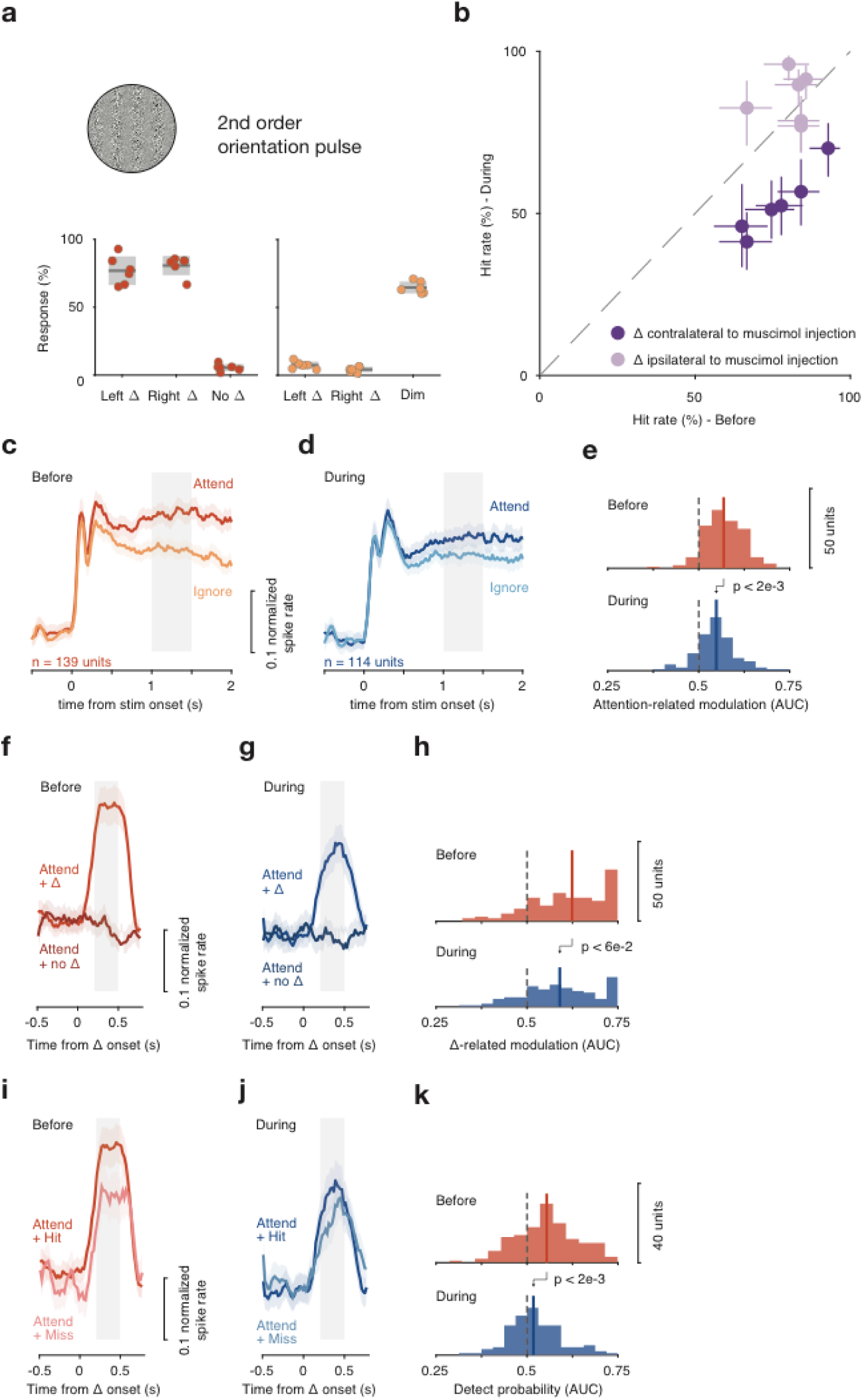
Effect of SC inactivation on attention-related modulation, change-related activity and detect probability in fSTS neurons was not specific to motion stimuli. a. A 2^nd^ order orientation pulse stimulus (that contains no motion energy) was used in 6 SC inactivation sessions to evaluate whether the effects of SC inactivation on fSTS neurons was specific to motion stimulus (see methods). Top: image of orientation pulse. Bottom: behavioral performance in the task (similar format to figure 1a, b). b. A comparison of hit rate (i.e. the successful detection of an orientation pulse) for contralateral and ipsilateral stimuli, in the *Attend* condition, before and during SC inactivation. The observed contralateral deficits with the second-order stimulus are consistent with a previous report ^8^. c-e. Attention-related modulation before and during SC inactivation (same format as figure 2a-c). f-h. Change-related activity before and during SC inactivation (same format as figure 3a-c). i-k. Detect probability before and during SC inactivation (same format as figure 3e-g).

**Supplementary Figure 9.**
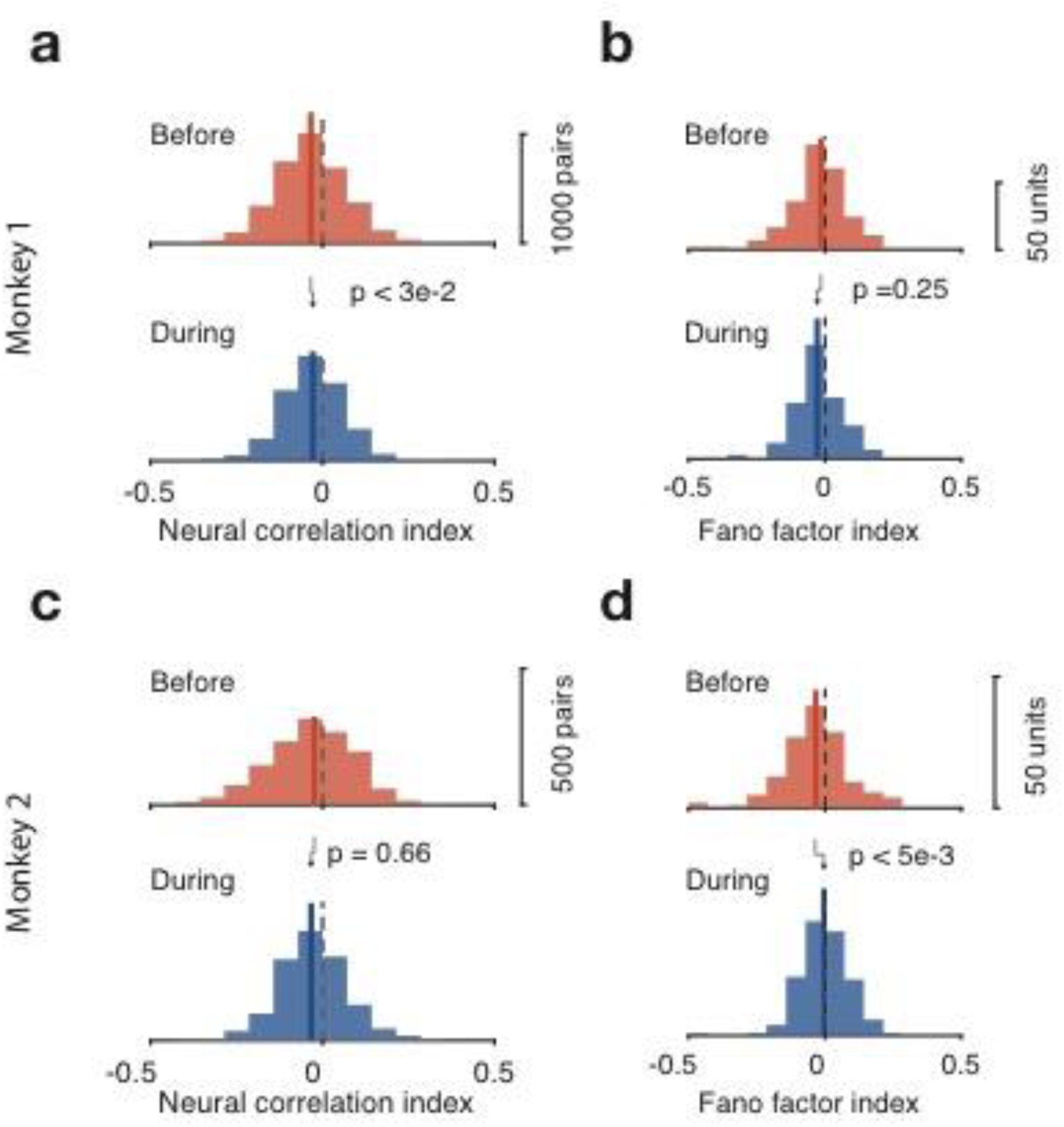
Effect of SC inactivation on spike count correlations and fano-factor in fSTS neurons. a. Distribution of neuronal correlation indices (see Methods) before (top) and during (bottom) SC inactivation, for monkey 1. b. Distribution of neuronal fano-factor indices (see Methods) before (top) and during (bottom) SC inactivation, for monkey 1. c. Same as a, for monkey 2. d. Same as b, for monkey 2.

**Supplementary Figure 10.**
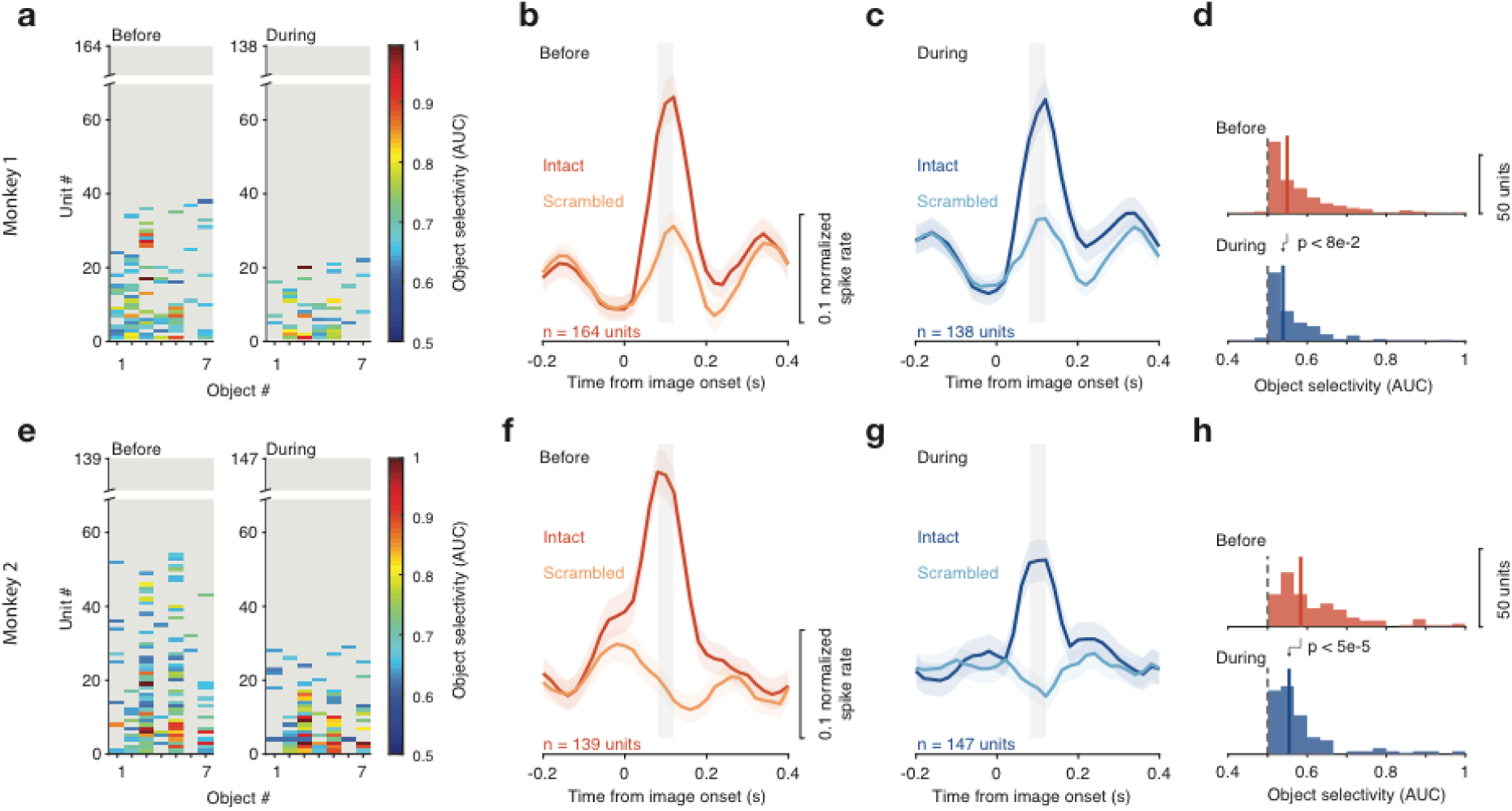
Effect of SC inactivation on object selectivity in fSTS neurons is similar across monkeys. a. Object selectivity across single neurons and the population, before and during SC inactivation, in monkey 1 (same format as figure 4c-f). b. Object selectivity across single neurons and the population, before and during SC inactivation, in monkey 2 (same format as figure 4c-f).

**Supplementary Figure 11.**
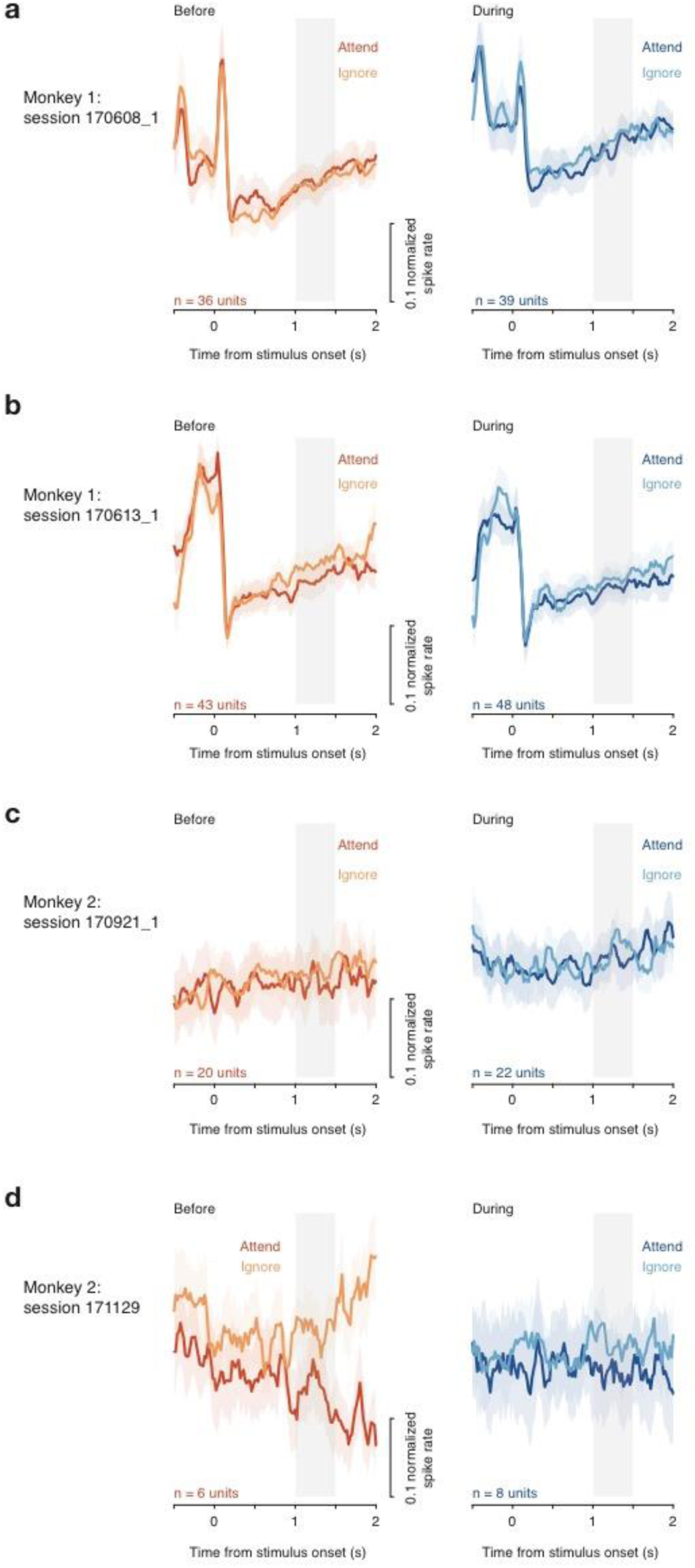
Neurons outside of the circumscribed fMRI ‘hotspot’ show non-standard responses. a-d. Population average of neuronal responses, recorded in locations outside of the putative hotspot (monkey 1: a, b; monkey 2: c, d), to motion stimulus in the *Attend* and *Ignore* conditions, before (left) and during (right) SC inactivation.

